# A *de novo* variant in *PAK2* detected in an individual with Knobloch type 2 syndrome

**DOI:** 10.1101/2024.04.18.590108

**Authors:** Elizabeth A. Werren, Louisa Kalsner, Jessica Ewald, Michael Peracchio, Cameron King, Purva Vats, Peter A. Audano, Peter N. Robinson, Mark D. Adams, Melissa A. Kelly, Adam P. Matson

## Abstract

P21-activated kinase 2 (PAK2) is a serine/threonine kinase essential for a variety of cellular processes including signal transduction, cellular survival, proliferation, and migration. A recent report proposed monoallelic *PAK2* variants cause Knobloch syndrome type 2 (KNO2)—a developmental disorder primarily characterized by ocular anomalies. Here, we identified a novel *de novo* heterozygous missense variant in *PAK2*, NM_002577.4:c.1273G>A, p.(D425N), by whole genome sequencing in an individual with features consistent with KNO2. Notable clinical phenotypes include global developmental delay, congenital retinal detachment, mild cerebral ventriculomegaly, hypotonia, FTT, pyloric stenosis, feeding intolerance, patent ductus arteriosus, and mild facial dysmorphism. The p.(D425N) variant lies within the protein kinase domain and is predicted to be functionally damaging by *in silico* analysis. Previous clinical genetic testing did not report this variant due to unknown relevance of *PAK2* variants at the time of testing, highlighting the importance of reanalysis. Our findings also substantiate the candidacy of *PAK2* variants in KNO2 and expand the KNO2 clinical spectrum.

## INTRODUCTION

Neurodevelopmental disorders (NDD) exhibit high genetic heterogeneity, often with non-specific syndromic features, which poses major challenges for molecular diagnosis. Knobloch syndrome (KNO) is a developmental disorder characterized by ocular abnormalities including high myopia, vitreoretinal degeneration, retinal detachment, and variable occipital skull defects. While many cases of KNO are caused by variants in *COL18A1*, diagnosed as Knobloch syndrome type 1 (KNO1, MIM:120328), other individuals with clinically diagnosed KNO have been identified who lack variants in this gene.^1^ Recently, *PAK2* was identified as a candidate gene for autosomal dominant Knobloch syndrome type 2 (KNO2; MIM:618458), which has similar features to KNO1.^1^

PAK2 is a member of the p21-activated kinase (PAK) family proteins, which are serine/threonine kinases involved in signal transduction, cytoskeletal dynamics, cell cycle progression, and cell motility.^2,3^ There are two subfamilies of PAK proteins based on structure, regulation, and sequence identity. Group I PAK proteins are activated by Rho GTPases CDC42 and RAC1, and include PAK1, PAK2, and PAK3—each of which has been implicated in NDD (MIM:618158, MIM:618458, and MIM:300558, respectively).^3^ Conversely, group II PAK proteins (PAK4, PAK5, and PAK6) are activated independently of Rho GTPases and have not been associated with NDD to date.^3^

*PAK2* is located within the chromosome 3q29 region that is commonly rearranged in NDD, wherein 3q29 duplications cause intellectual disability (ID), microcephaly, cerebral palsy, and epilepsy, and 3q29 deletions cause autism spectrum disorder (ASD).^4,5^ Aside from a missense variant in *PAK2* detected in individuals with KNO2^1^, two missense and one nonsense *PAK2* variants have been implicated in ASD.^6^ Given the paucity of reported *PAK2* variants in NDD, our understanding of *PAK2* genotype-phenotype correlations is limited.

Here, we performed whole genome sequencing (WGS) on an individual with congenital retinal detachment, hypotonia, global developmental delay (GDD), mild cerebral ventriculomegaly, failure to thrive (FTT) with pyloric stenosis and persistent feeding intolerance, patent ductus arteriosus (PDA) requiring repair, and mild dysmorphisms. Prior clinical testing, including exome sequencing and chromosomal microarray, was negative (GeneDx). Using short-read (SR-) and long-read (LR-) WGS, a novel *de novo* heterozygous variant was discovered in *PAK2* (c.1273G>A, p.(D425N)). Given the phenotypic overlap with previously published KNO2 cases, we propose *PAK2* c.1273G>A, p.(D425N) as a candidate pathogenic variant in KNO2. This case report expands the phenotypic and genotypic spectra of *PAK2*-associated KNO2.

## MATERIALS AND METHODS

### Participant recruitment and sample collection

Participants provided informed consent in accordance with the ethical standards for human research subjects of the IRB committees at the Connecticut Children’s Medical Center and The Jackson Laboratory for Genomic Medicine. Venous whole blood samples were obtained from consented subjects using standard collection procedures and EDTA tubes.

### Genome sequencing and analysis

Research trio LR-WGS and singleton SR-WGS were performed at The Jackson Laboratory for Genomic Medicine. Variant confirmation was performed by review of prior exome sequencing at GeneDx.

Genomic DNA (gDNA) was extracted from whole blood using the Qiagen PAXgene Blood DNA kit (Cat. #: 761133). For LR-WGS, BluePippin size selection was performed, and samples were prepped using the PacBio SMRTBell prep kit 3.0 and SMRTbell barcoded adapters. The libraries were assessed for quality and sequenced using a PacBio Revio instrument. Mean coverage was 37.5X, 34.5X, and 37.9X for the proband, mother, and father, respectively. For SR-WGS, proband gDNA was fragmented and ligated to unique index adapters using an Illumina DNA PCR-free library preparation. The library was assessed for quality and sequenced using an Illumina NovaSeq 6000 instrument (150-bp paired-end reads) to 47.3X mean coverage.

LR-WGS analysis was performed using custom, in-house secondary and tertiary analysis pipelines with phased-assembly and read-based analysis workflows. For the *de novo* assembly-based pipeline, trio-informed phased-assembly was performed using Hifiasm with Trio binning (v0.19.7-r598), followed by variant calling with PAV (v2.3.4) and variant prioritization by Human Phenotype Ontology (HPO) terms using Exomiser (v13.3.0).^7-9^ For PAV, calls were made from the phased assembly against the no-alternative (no-ALT) GRCh38 reference genome assembly by Human Genome Structural Variation (HGSV) (ftp://ftp.1000genomes.ebi.ac.uk/vol1/ftp/data_collections/HGSVC2/technical/reference/20200513_hg38_NoALT/). For the alignment-based workflow, pbmm2 (v1.13.1) was used to align reads to the no-ALT GRCh38 reference. Read-based phasing was then performed using DeepVariant (v1.6.0) and WhatsHap (v1.7).^10,11^ Small variant calling was performed with Deepvariant (v1.6.0).^10^ Structural variants were called with PacBio pbsv (v2.9.0) and Sniffles2 (v2.0.7).^12^ Variant prioritization by HPO terms was performed using Exomiser (v13.3.0) and SvAnna (v1.0.4).^8,13^ For SR-WGS processing, output sequencing data were converted from BCL to FASTQ format. Alignment to the human reference genome GRCh38, variant calling, and annotation were performed using the Illumina Emedgene platform (v33.0.17).

Phenotype prioritization for both LR-WGS and SR-WGS pipelines was performed using the following HPO terms: HP:0001263, GDD; HP:0000541, retinal detachment; HP:0008936, axial hypotonia; HP:0002119, ventriculomegaly; HP:0002021, pyloric stenosis; HP:0001508, FTT; HP:0033454, tube feeding; HP:0001643, patent ductus arteriosus. HPO terms for the present study are also provided as a phenopacket (Supplementary File 1). Variants were interpreted using ACMG guidelines.^14^

### Protein conservation, structure, and in silico analyses

NCBI HomoloGene tool (https://www.ncbi.nlm.nih.gov/homologene) was used to obtain aligned amino acid sequences of PAK2 across species at affected residues and flanking regions. The PDB model file for the PAK2 kinase domain (6FD3_A) was downloaded and the structure was edited in PyMOL v2.5.2. *In silico* prediction of the functional impact of the *PAK2* variant was performed with PolyPhen-2 (v2.2.3r406) using the HumDiv model (http://genetics.bwh.harvard.edu/pph2), SIFT (https://sift.bii.a-star.edu.sg/), and Missense3D-Database (http://missense3d.bc.ic.ac.uk:8080).

Combined Annotation Dependent Depletion (CADD) Phred scores were obtained using CADD v1.6 against GRCh38 (https://cadd.gs.washington.edu). MetaDome analysis was performed on NM_002577.4 transcript using the online tool (stuart.radboudumc.nl/metadome).

## RESULTS

### Deleterious heterozygous variant in *PAK2* identified in an individual with NDD

The affected individual was a fifteen-month-old male who presented with a constellation of developmental features (Table 1; Supplementary File 1). He was delivered by cesarean section at 38.5 weeks’ gestation to a 24-year-old G1 mother. Both parents were unaffected (Figure 1A). Birth weight was 5 pounds, 12 ounces. He developed respiratory distress in the first days of life that required a one-month hospital stay in the neonatal intensive care unit. A large PDA was closed via cardiac catheter approach at 2 weeks of age. Due to migration of the device, he required repeat PDA closure at 3 months of age. He had persistent vomiting and poor weight gain. Pyloric stenosis was diagnosed and repaired at 3 months of age. A gastrostomy tube was placed, but he had persistent feeding intolerance with slow weight gain. A jejunostomy tube was inserted for continuous feeds with subsequent improvement in weight gain. At 15 months of age, weight and length were near the 50^th^ percentile, but persistent microcephaly was noted with a head circumference at the 1^st^ percentile.

**Table 1.**
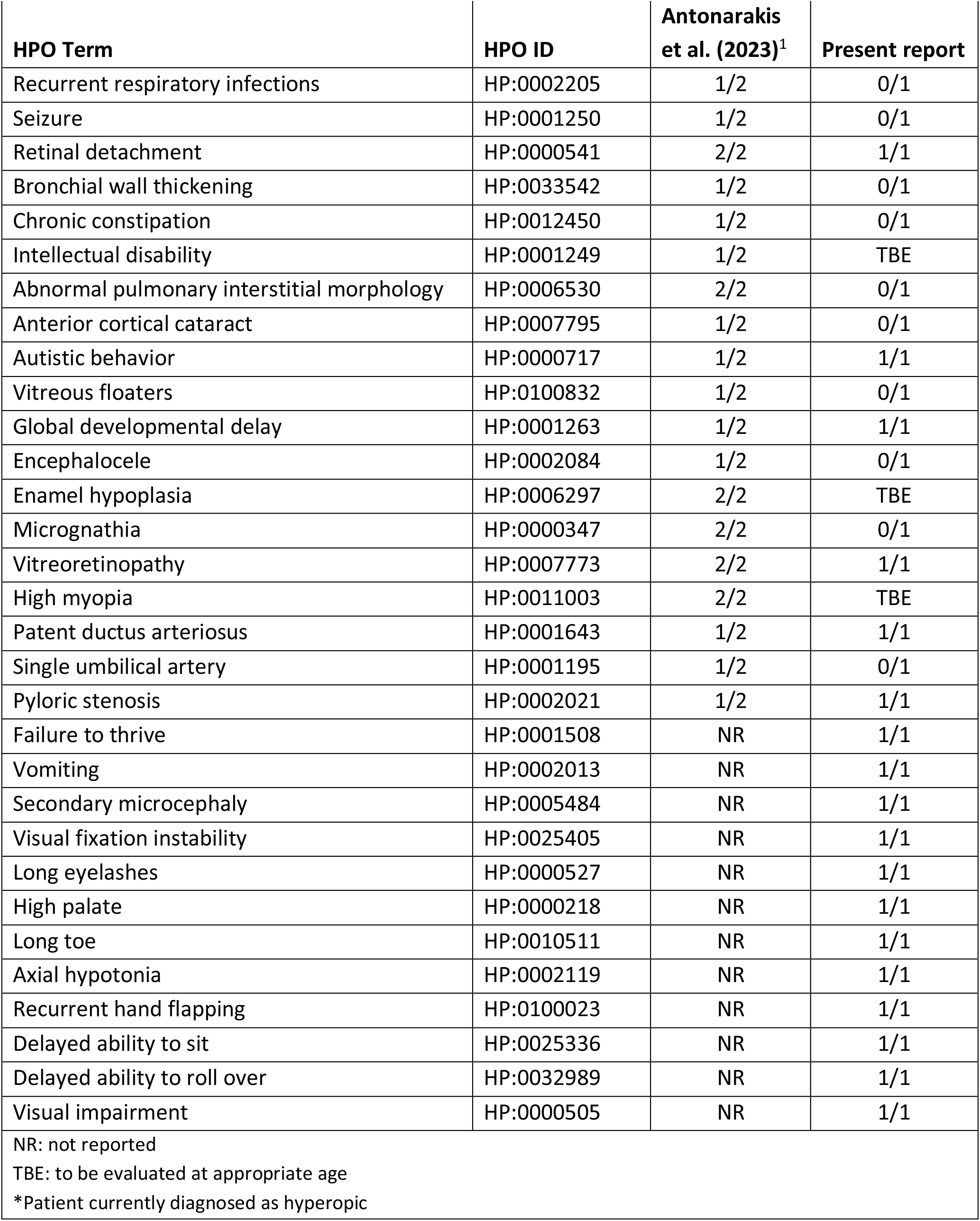
Comparison of clinical phenotypes observed in individuals with Knobloch syndrome 2.

**Figure 1.**
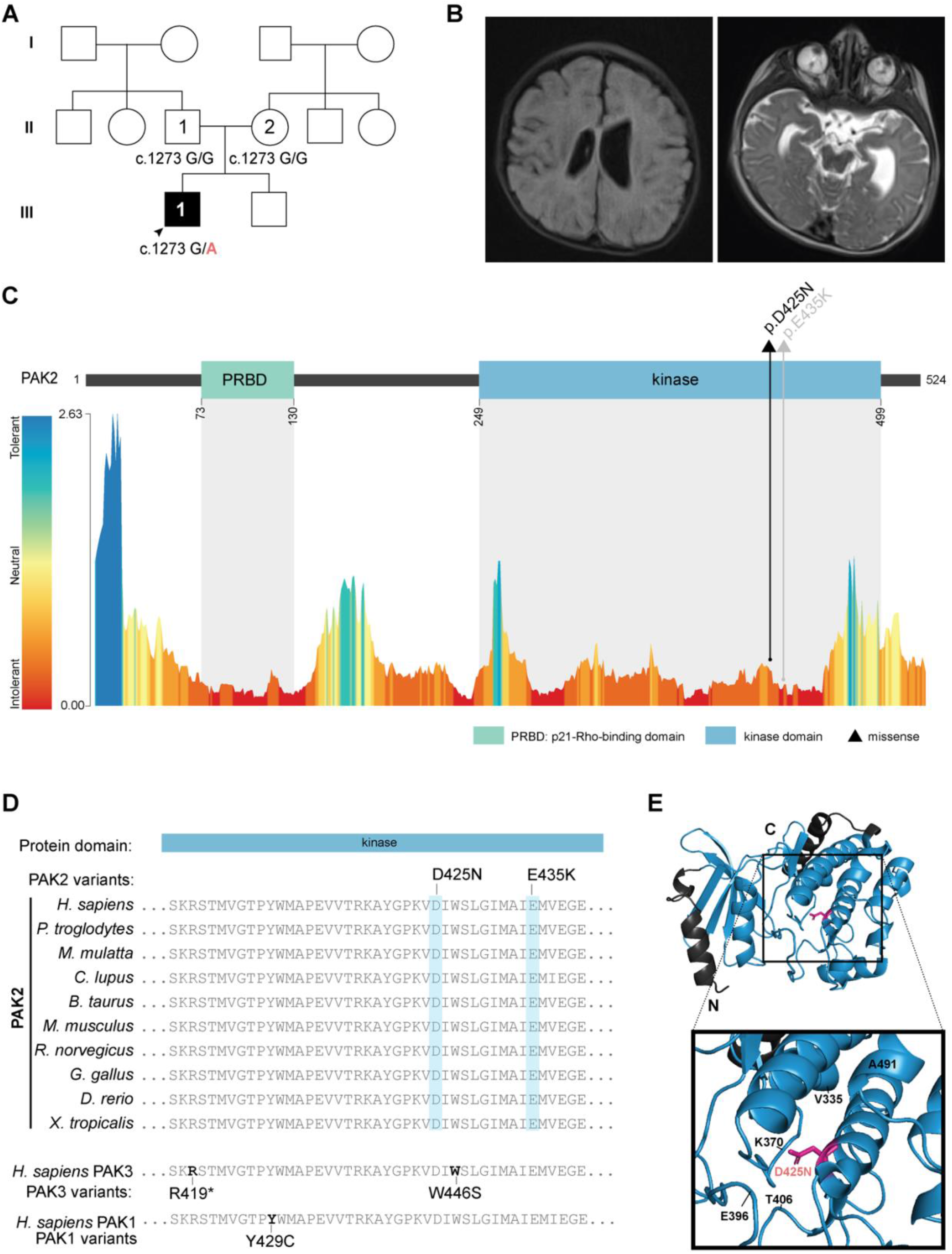
Monoallelic variants in *PAK2*. (**A**) Pedigree of the family in the present study with the genotypes for NM_002577.4:c.1273G>A (NC_000003.12:g.196820490G>A) of sequenced individuals. Squares represent assigned male at birth; circles represent assigned female at birth; clear shapes indicate unaffected status; solid shapes indicate affected status. (**B**) Cranial MRI demonstrating prominence of ventricles and sulci with asymmetric fullness of the body of the left ventricle as well as abnormal bilateral signal within the ocular globes. (**C**) Top: linear PAK2 protein map with missense variants (represented as a triangle) identified in the present study (black) and previously published (grey) in individuals with KNO2.^1^ PRBD, p21-Rho-binding domain (green) and protein kinase domain (blue). Bottom: predicted tolerance landscape of PAK2 (ENST00000327134.3) by MetaDome analysis. Tolerance scores are based on *dN/dS*. (**D**) PAK2 protein alignment showing cross-species conservation at affected residues, as well as protein alignment with the homologous kinase region in paralogs PAK3 and PAK1 (bottom) with homologous PAK3/PAK1 kinase variants. (**E**) PyMOL visualization of PAK2 protein kinase domain structure (PDB: 6FD3_A) showing the p.(D425N) variant.

At 4 months of age, he lacked visual fixation and tracking. Brain MRI revealed prominent ventricles and abnormal signals within the globes of both eyes (Figure 1B). Ophthalmological evaluation led to the diagnosis of complete left-sided retinal detachment with subretinal fibrosis and partial retinal detachment of the right eye. He underwent surgical repair of the retina at 7 months of age but has persistent visual impairment and wears corrective lenses. He had distinctive features including long eyelashes, high arched palate, and long toes. Neurological examination was notable for mild hypotonia and repetitive behaviors including frequent hand clapping. At 15 months of age, he was able to briefly sit unsupported but did not roll or stand.

Using both SR- and LR-WGS, a novel, *de novo* heterozygous missense variant in *PAK2* (NC_000003.12:g.196820490G>A, NM_002577.4:c.1273G>A, p.(D425N)) was discovered and confirmed by GeneDx by review of the prior exome sequencing. PAK2 contains two protein domains: a p21-Rho-binding domain (PRBD) and a protein kinase domain (Figure 1C). The PRBD domain interacts with active GTP-loaded RAC1 and CDC42, and the kinase domain affords PAK2 its catalytic function in phosphorylation.^15^ PAK2 is intolerant to both missense (Z = 1.83) and predicted loss-of-function (LOF; pLI = 0.83) variants in the general population (gnomADv4.0.0). The p.(D425N) missense variant lies within the protein kinase domain of PAK2, 10 residues upstream of a previously published KNO2 variant (p.E435K; Figure 1C).^1^ The p.(D425N) variant is absent in the general population (gnomADv4.0.0). It affects a residue that is intolerant to variation (*dN/dS* = 0.34, MetaDome, Figure 1C) and highly conserved across species (Figure 1D). PAK2 retains very high sequence homology with its paralogs, PAK1 and PAK3, at the protein kinase domain, and PAK2 missense variants map to nearby homologous residues affected in PAK1- and PAK3-associated developmental disorders (Figure 1D). *In silico* analysis predicts a deleterious effect of p.(D425N) on PAK2 protein (CADD Phred score: 31; PolyPhen2 HDivPred score: 1; SIFT score: 0.00). PAK2 p.(D425N) is predicted to be structurally damaging by Missense3D. Specifically, this substitution replaces a buried charged residue with an uncharged residue which is predicted to break a buried salt bridge (Figure 1E). Together, these findings suggest a pathogenic effect of the PAK2 p.(D425N) variant (ClinVar Variation ID: 2921281).

## DISCUSSION

Knobloch syndrome is characterized by retinal detachment and occipital encephalocele. The more common KNO1 is associated with cognitive impairment and epilepsy (MIM:267750). KNO2, thus far, has been reported in few individuals and manifests with ocular involvement, scalp or skull defects, dysmorphic features, pyloric stenosis, PDA, abnormal interstitial markings in the lung, structural brain defects, epilepsy, and cognitive impairment (MIM:618458). Prior genetic testing approaches did not reveal any candidate pathogenic variants in genes that could explain the full clinical presentation of the affected individual in this study. Using SR- and LR-WGS, we identified a novel *de novo* heterozygous dominant variant in *PAK2* as a candidate genetic variant in KNO2. In a previous report, two affected siblings harbored the *PAK2* c.1303G>A, p.(E435K) variant, likely inherited via germline mosaicism due to absence in the parents.^1^ Both brothers displayed severe retinal detachment, a characteristic phenotype shared with the present case (Table 1).^1^ The younger brother had encephalocele and interstitial lung changes on X-ray with frequent respiratory infection and constipation, phenotypes not observed in the present study, demonstrating the variability in KNO2 (Table 1).^1^

Shared phenotypes in both KNO2 and the other PAK-associated disorders include GDD, ID, ASD, and epilepsy, though these findings may be variable. In addition, variants in *PAK1* are also associated with macrocephaly whereas those in *PAK3* are associated with microcephaly, a phenotype observed in the present report. Other recurring features in PAK-related disorders include hypotonia, speech difficulties, neurological/neuromuscular defects, skin abnormalities, and facial dysmorphisms. Of these, hypotonia and dysmorphic features (high arched palate, long eyelashes, and long toes) are also noted in the individual in the present study. Previous work in *C. elegans* has demonstrated functional redundancy of PAK2 and PAK1 in the regulation of cellular migration, indicating a possibility of compensatory mechanisms at play in affected cells.^16^ While group I PAK proteins retain structural and regulatory similarity with overlapping substrates, their expression profiles vary (GTEx Portal, BrainSpan Atlas), which may contribute to the phenotypic differences observed between PAK disorders.^17^

Nearly all pathogenic variants reported in group I *PAK* genes map to the PRBD or kinase domains, correlating to the regions with the highest variation intolerance across the protein (Figure 1C). Among these variants is a greater portion of missense variants that map to the PAK protein kinase domains (partly shown in Figure 1D). The majority of PAK kinase variants have been characterized to be inactivating as measured by disrupted kinase activity; however, a handful result in gain-of-function (GOF).^1,18-20^ Moreover, the genetic mechanisms of pathogenic variants differ across PAK proteins, with *PAK1* variants exerting a dominant GOF effect, as measured by elevated *PAK1* activity in patient cells.^18^ Conversely, the previously published PAK2 p.E435K missense variant results in PAK2 haploinsufficiency.^1^ In the case of *PAK3*, both LOF and GOF variants have been described.^18-20^ *In silico* analyses and structural modeling suggest a deleterious effect of PAK2 p.(D425N), and in vivo studies for the p.E435K variant demonstrate reduced activity; however, further investigation will be required to determine the functional impact of the p.(D425N) variant.

In summary, we report one individual with a *de novo* heterozygous variant in *PAK2* and NDD characterized by GDD, retinal detachment, PDA, pyloric stenosis, prominent cerebral ventricles, and mild dysmorphism. This case report substantiates PAK2’s role in KNO2 and expands the genetic and phenotypic landscape of the disease.

## ACKNOWLEDGEMENTS

We would like to thank the family for their participation in this work. We would like to thank Charles Lee, Juan C. Salazar, Christine R. Beck, Alyx Vogle, Elizabeth Charnysh, and Kunal Sanghavi for their intellectual contributions. We would like to thank GeneDx for use of data. This work was supported by funds from the Connecticut Children’s Research Institute and the Jackson Laboratory for Genomic Medicine. P.A.A was supported by NIH NIGMS R35GM133600 and NIH NCI P30CA034196.

## REFERENCES

1. Antonarakis SE, Holoubek A, Rapti M, et al. Dominant monoallelic variant in the PAK2 gene causes Knobloch syndrome type 2. Hum Mol Genet 2021;31(1):1–9. doi: 10.1093/hmg/ddab026

2. Rudel T, Bokoch GM. Membrane and morphological changes in apoptotic cells regulated by caspase-mediated activation of PAK2. Science 1997;276(5318):1571–4. doi: 10.1126/science.276.5318.1571

3. Eswaran J, Soundararajan M, Kumar R, et al. UnPAKing the class differences among p21-activated kinases. Trends Biochem Sci 2008;33(8):394–403. doi: 10.1016/j.tibs.2008.06.002 [published Online First: 20080717]

4. Sanders SJ, He X, Willsey AJ, et al. Insights into Autism Spectrum Disorder Genomic Architecture and Biology from 71 Risk Loci. Neuron 2015;87(6):1215–33. doi: 10.1016/j.neuron.2015.09.016 [published Online First: 2015/09/25]

5. Wen X, Zhu J, Cai L, et al. A familial 3q28q29 duplication induced mild intellectual disability: case presentation and literature review. Am J Transl Res 2022;14(3):1663–71. [published Online First: 20220315]

6. Wang Y, Zeng C, Li J, et al. PAK2 Haploinsufficiency Results in Synaptic Cytoskeleton Impairment and Autism-Related Behavior. Cell Rep 2018;24(8):2029–41. doi: 10.1016/j.celrep.2018.07.061

7. Ebert P, Audano PA, Zhu Q, et al. Haplotype-resolved diverse human genomes and integrated analysis of structural variation. Science 2021;372(6537) doi: 10.1126/science.abf7117 [published Online First: 20210225]

8. Robinson PN, Kohler S, Oellrich A, et al. Improved exome prioritization of disease genes through cross-species phenotype comparison. Genome Res 2014;24(2):340–8. doi: 10.1101/gr.160325.113

9. Cheng H, Concepcion GT, Feng X, et al. Haplotype-resolved de novo assembly using phased assembly graphs with hifiasm. Nat Methods 2021;18(2):170–75. doi: 10.1038/s41592-020-01056-5 [published Online First: 20210201]

10. Poplin R, Chang PC, Alexander D, et al. A universal SNP and small-indel variant caller using deep neural networks. Nat Biotechnol 2018;36(10):983–87. doi: 10.1038/nbt.4235 [published Online First: 20180924]

11. Martin M, Ebert P, Marschall T. Read-Based Phasing and Analysis of Phased Variants with WhatsHap. Methods Mol Biol 2023;2590:127–38. doi: 10.1007/978-1-0716-2819-5_8

12. Smolka M, Paulin LF, Grochowski CM, et al. Detection of mosaic and population-level structural variants with Sniffles2. Nat Biotechnol 2024 doi: 10.1038/s41587-023-02024-y [published Online First: 20240102]

13. Danis D, Jacobsen JOB, Balachandran P, et al. SvAnna: efficient and accurate pathogenicity prediction of coding and regulatory structural variants in long-read genome sequencing. Genome Med 2022;14(1):44. doi: 10.1186/s13073-022-01046-6 [published Online First: 20220428]

14. Richards S, Aziz N, Bale S, et al. Standards and guidelines for the interpretation of sequence variants: a joint consensus recommendation of the American College of Medical Genetics and Genomics and the Association for Molecular Pathology. Genet Med 2015;17(5):405–24. doi: 10.1038/gim.2015.30 [published Online First: 2015/03/05]

15. Lei M, Lu W, Meng W, et al. Structure of PAK1 in an autoinhibited conformation reveals a multistage activation switch. Cell 2000;102(3):387–97. doi: 10.1016/s0092-8674(00)00043-x

16. Peters EC, Gossett AJ, Goldstein B, et al. Redundant canonical and noncanonical Caenorhabditis elegans p21-activated kinase signaling governs distal tip cell migrations. G3 (Bethesda) 2013;3(2):181–95. doi: 10.1534/g3.112.004416 [published Online First: 20130201]

17. Wang Y, Guo F. Group I PAKs in myelin formation and repair of the central nervous system: what, when, and how. Biol Rev Camb Philos Soc 2022;97(2):615–39. doi: 10.1111/brv.12815 [published Online First: 20211122]

18. Harms FL, Kloth K, Bley A, et al. Activating Mutations in PAK1, Encoding p21-Activated Kinase 1, Cause a Neurodevelopmental Disorder. Am J Hum Genet 2018;103(4):579–91. doi: 10.1016/j.ajhg.2018.09.005

19. Duarte K, Heide S, Poëa-Guyon S, et al. PAK3 mutations responsible for severe intellectual disability and callosal agenesis inhibit cell migration. Neurobiol Dis 2020;136:104709. doi: 10.1016/j.nbd.2019.104709 [published Online First: 20191214]

20. Hertecant J, Komara M, Nagi A, et al. A de novo mutation in the X-linked PAK3 gene is the underlying cause of intellectual disability and macrocephaly in monozygotic twins. Eur J Med Genet 2017;60(4):212–16. doi: 10.1016/j.ejmg.2017.01.004 [published Online First: 20170124]

