## Supplemental File 1 for "A *de novo* variant in *PAK2* detected in an individual with Knobloch type 2 syndrome"

```
{
  "id": "affected_individual_",
  "subject": {
    "id": "affected individual ",
    "timeAtLastEncounter": {
      "age": {
        "iso8601duration": "P1Y3M"
      }
    },
    "sex": "MALE"
  },
  "phenotypicFeatures": [
    {
      "type": {
        "id": "HP:0000541",
        "label": "Retinal detachment"
      },
      "onset": {
        "age": {
          "iso8601duration": "P1D"
        }
      }
    },
    {
      "type": {
        "id": "HP:0001643",
        "label": "Patent ductus arteriosus"
      },
      "onset": {
        "age": {
          "iso8601duration": "P1D"
        }
      }
    },
    {
      "type": {
        "id": "HP:0001508",
        "label": "Failure to thrive"
      },
      "onset": {
        "age": {
          "iso8601duration": "P3M"
        }
      }
    },
    {
      "type": {
        "id": "HP:0001643",
        "label": "Tube feeding"
      },
      "onset": {
        "age": {
          "iso8601duration": "P3M"
        }
      }
    },
```

```
{
  "type": {
    "id": "HP:0002021",
    "label": "Pyloric stenosis"
  },
  "onset": {
    "age": {
      "iso8601duration": "P3M"
    }
  }
},
{
  "type": {
    "id": "HP:0002013",
    "label": "Vomiting"
  },
  "onset": {
    "age": {
      "iso8601duration": "P3M"
    }
  }
},
{
  "type": {
    "id": "HP:0005484",
    "label": "Secondary microcephaly"
  },
  "onset": {
    "age": {
      "iso8601duration": "P1Y3M"
    }
  }
},
{
  "type": {
    "id": "HP:0025405",
    "label": "Visual fixation instability"
  },
  "onset": {
    "age": {
      "iso8601duration": "P4M"
    }
  }
},
{
  "type": {
    "id": "HP:0000541",
    "label": "Retinal detachment"
  },
  "onset": {
    "age": {
      "iso8601duration": "P4M"
    }
  }
},
{
```

```
      "type": {
        "id": "HP:0002021",
        "label": "Ventriculomegaly"
      },
      "onset": {
        "age": {
          "iso8601duration": "P4M"
        }
      }
    },
    {
      "type": {
        "id": "HP:0000527",
        "label": "Long eyelashes"
      },
      "onset": {
        "age": {
          "iso8601duration": "P7M"
        }
      }
    },
    {
      "type": {
        "id": "HP:0000218",
        "label": "High palate"
      },
      "onset": {
        "age": {
          "iso8601duration": "P7M"
        }
      }
    },
    {
      "type": {
        "id": "HP:0010511",
        "label": "Long toe"
      },
      "onset": {
        "age": {
          "iso8601duration": "P7M"
        }
      }
    },
    {
      "type": {
        "id": "HP:0002119",
        "label": "Axial hypotonia"
      },
      "onset": {
        "age": {
          "iso8601duration": "P7M"
        }
      }
    },
    {
      "type": {
```

```
        "id": "HP:0100023",
        "label": "Recurrent hand flapping"
      },
      "onset": {
        "age": {
          "iso8601duration": "P7M"
        }
      }
    },
    {
      "type": {
        "id": "HP:0025336",
        "label": "Delayed ability to sit"
      },
      "onset": {
        "age": {
          "iso8601duration": "P7M"
        }
      }
    },
    {
      "type": {
        "id": "HP:0032989",
        "label": "Delayed ability to roll over"
      },
      "onset": {
        "age": {
          "iso8601duration": "P7M"
        }
      }
    },
    {
      "type": {
        "id": "HP:0000505",
        "label": "Visual impairment"
      },
      "onset": {
        "age": {
          "iso8601duration": "P7M"
        }
      }
    },
    {
      "type": {
        "id": "HP:0001263",
        "label": "Global developmental delay"
      },
      "onset": {
        "age": {
          "iso8601duration": "P1Y3M"
        }
      }
    }
  ],
  "interpretations": [
    {
```

```
    "id": "affected individual ",
    "progressStatus": "SOLVED",
    "diagnosis": {
      "disease": {
        "id": "OMIM:618458",
        "label": "Knobloch syndrome 2"
      },
      "genomicInterpretations": [
        {
          "subjectOrBiosampleId": "affected individual ",
          "interpretationStatus": "CAUSATIVE",
          "variantInterpretation": {
            "variationDescriptor": {
              "id": "var_hWhSlQleEoWlPsljPhhwcAkmR",
              "geneContext": {
                "valueId": "HGNC:8591",
                "symbol": "PAK2"
              },
              "expressions": [
                {
                  "syntax": "hgvs.c",
                  "value": "NM_002577.4:c.1273G>A"
                },
                {
                  "syntax": "hgvs.g",
                  "value": "NC_000003.12:g.196820490G>A"
                }
              ],
              "vcfRecord": {
                "genomeAssembly": "hg38",
                "chrom": "chr3",
                "pos": "196820490",
                "ref": "G",
                "alt": "A"
              },
              "moleculeContext": "genomic",
              "allelicState": {
                "id": "GENO:0000135",
                "label": "heterozygous"
              }
            }
          }
        }
      ]
    }
  }
],
"metaData": {
  "created": "2024-01-20T15:02:29.990556955Z",
  "createdBy": "ORCID:0000-0002-0736-9199",
  "resources": [
    {
      "id": "geno",
      "name": "Genotype Ontology",
      "url": "http://purl.obolibrary.org/obo/geno.owl",
      "version": "2022-03-05",
```

```
        "namespacePrefix": "GENO",
        "iriPrefix": "http://purl.obolibrary.org/obo/GENO_"
      },
      {
        "id": "hgnc",
        "name": "HUGO Gene Nomenclature Committee",
        "url": "https://www.genenames.org",
        "version": "06/01/23",
        "namespacePrefix": "HGNC",
        "iriPrefix": "https://www.genenames.org/data/gene-symbol-report/#!/hgnc_id/"
      },
      {
        "id": "omim",
        "name": "An Online Catalog of Human Genes and Genetic Disorders",
        "url": "https://www.omim.org",
        "version": "January 4, 2023",
        "namespacePrefix": "OMIM",
        "iriPrefix": "https://www.omim.org/entry/"
      },
      {
        "id": "so",
        "name": "Sequence types and features ontology",
        "url": "http://purl.obolibrary.org/obo/so.obo",
        "version": "2021-11-22",
        "namespacePrefix": "SO",
        "iriPrefix": "http://purl.obolibrary.org/obo/SO_"
      },
      {
        "id": "hp",
        "name": "human phenotype ontology",
        "url": "http://purl.obolibrary.org/obo/hp.owl",
        "version": "2024-01-16",
        "namespacePrefix": "HP",
        "iriPrefix": "http://purl.obolibrary.org/obo/HP_"
      }
    ],
    "phenopacketSchemaVersion": "2.0"
  }
}
```
